# Spatial modeling of prostate cancer metabolic gene expression reveals extensive heterogeneity and selective vulnerabilities

**DOI:** 10.1101/719294

**Authors:** Yuliang Wang, Shuyi Ma, Walter L. Ruzzo

**Affiliations:** Institute for Stem Cell and Regenerative Medicine, University of Washington, Seattle, WA 98109, USA; Paul G. Allen School of Computer Science & Engineering, University of Washington, Seattle, WA 98195, USA; Center for Global Infectious Disease Research, Seattle Children’s Research Institute, Seattle WA, USA; Department of Genome Sciences, University of Washington School of Medicine, Seattle, WA, 98195, USA; Fred Hutchinson Cancer Research Center, Seattle, WA, 98102, USA

**Author notes:** Correspondence should be addressed to Y.W.

## Abstract

Spatial heterogeneity is a fundamental feature of the tumor microenvironment (TME), and tackling spatial heterogeneity in neoplastic metabolic aberrations is critical for tumor treatment. Genome-scale metabolic network models have been used successfully to simulate cancer metabolic networks. However, most models use bulk gene expression data of entire tumor biopsies, ignoring spatial heterogeneity in the TME. To account for spatial heterogeneity, we performed spatially-resolved metabolic network modeling of the prostate cancer microenvironment. We discovered novel malignant-cell-specific metabolic vulnerabilities targetable by small molecule compounds. We predicted that inhibiting the fatty acid desaturase SCD1 may selectively kill cancer cells based on our discovery of spatial separation of fatty acid synthesis and desaturation. We also uncovered higher prostaglandin metabolic gene expression in the tumor, relative to the surrounding tissue. Therefore, we predicted that inhibiting the prostaglandin transporter SLCO2A1 may selectively kill cancer cells. Importantly, SCD1 and SLCO2A1 have been previously shown to be potently and selectively inhibited by compounds such as CAY10566 and suramin, respectively. We also uncovered cancer-selective metabolic liabilities in central carbon, amino acid, and lipid metabolism. Our novel cancer-specific predictions provide new opportunities to develop selective drug targets for prostate cancer and other cancers where spatial transcriptomics datasets are available.

## Introduction

Cancer cells reprogram their metabolism to fulfill the energetic and biosynthetic needs of proliferation, invasion and migration^1^. This is exemplified in prostate cancer, the second most common cancer in American men after melanoma^2^. Previous studies have uncovered profound metabolic dysregulation in multiple pathways, particularly in fatty acid and lipid metabolism^3, 4^. Discovering novel cancer-specific metabolic aberrations has significant translational applications, because cancer-associated metabolic dysfunctions can be exploited to advance cancer detection (e.g., ^18^F-FDG (Fludeoxyglucose) imaging based on elevated glycolysis in cancer^5^) and treatment (e.g., L-asparaginase in treating acute lymphoblastic leukemia^6^).

Cancer metabolic reprograming is profoundly impacted by spatial heterogeneity, a fundamental feature of the tumor microenvironment (TME)^7^. Heterogeneous distributions of blood vessels and stromal tissues create uneven spatial gradients of nutrients and metabolic byproducts, which significantly shape the phenotypes of many cell types in the TME^8^. Recent technologies, such as spatial transcriptomics^9, 10^ and Slide-seq^11^ have enabled transcriptomic profiling of hundreds of locations within tissue sections with high spatial resolution (2-100 μm), and have been used to study multiple types of malignancies, including prostate cancer, breast cancer, pancreatic cancer, and melanoma^9, 10, 12–14^. These spatially-resolved datasets provide novel opportunities to dissect spatial metabolic heterogeneity in the TME and uncover novel tumor-specific metabolic vulnerabilities. However, due to the complexity of the cancer metabolic landscape^15^, uncovering the mechanistic connections of many spatially heterogeneous metabolic enzymes and evaluating their effects on cancer proliferation has been a significant challenge.

Genome-scale metabolic models (GEMs) are a computational framework that connect the thousands of metabolic enzymes, transporters and metabolites into a computable model. GEMs enable systematic *in silico* simulation of how metabolic perturbations affect cellular phenotypes such as growth and energy production. GEMs have been used to develop new strategies to selectively target cancer metabolism^16, 17^, including in prostate cancer^18^. However, current cancer GEMs are mostly based on bulk transcriptomics data that do not capture the spatial or cellular heterogeneity of the tumor microenvironment (TME).

To characterize cancer-specific metabolic vulnerabilities, we have developed a novel pipeline to build spatially resolved metabolic network models for prostate cancer using publicly available spatial transcriptomics data^12^. We identified metabolic genes and pathways with distinct spatial expression patterns that differ across separate tissue sections of the same primary tumor. This suggests that under a set of common hallmarks of cancer metabolism, tumor cells develop diverse survival strategies adapted to their local microenvironments. We also found malignant-cell-specific metabolic vulnerabilities by systematic *in silico* simulation, many of which have strong literature support. These genes can be targeted by potent and selective small molecule chemical compounds, some of which are already FDA-approved. This study demonstrated that spatially-resolved metabolic network models can generate mechanistic and clinically relevant insights into the metabolic complexities in the TME. The computational approach developed in this study represents an important first step to understand and untangle spatial metabolic heterogeneity. As spatial transcriptomics becomes increasingly used to characterize molecular heterogeneity in the tumor microenvironment of multiple types of cancer^9, 10, 12–14^, we expect that our novel modeling pipeline will provide a useful tool set to inform contextualization and interpretation of these complex datasets.

## Results

### Intra-tumor heterogeneity of spatially variable metabolic genes and pathways

We focused our analysis on previously published spatial transcriptomics data for three tumor tissue sections (numbered 1.2, 2.4 and 3.3) from the same primary tumor of a prostate cancer patient^12^. Transcriptome-wide data (3000 expressed genes per location on average) were available for hundreds of locations within each of the three tissue sections. The regions as outlined in Berglund *et al*^12^ with malignant cells circled as in (Figure 1A). These outlines were inferred from spatial transcriptomics data using a factor analysis method and confirmed by immunohistochemical staining^12^. We identified spatially variable (SV) genes using the spatialDE method^19^. SV genes show differential expression that significantly co-varies with spatial coordinates (i.e., adjacent locations have similar expression levels but distal locations have different expression). Figure 1B shows two examples. ACSL5 is spatially variable in tissue section 1.2 (highly expressed mainly in the tumor), while LRP1 is not (erratically expressed across the entire section). Compared to the analysis done in the Berglund *et al* study, spatialDE was more tailored for identifying specific genes with significant spatial variation.

**Figure 1.**
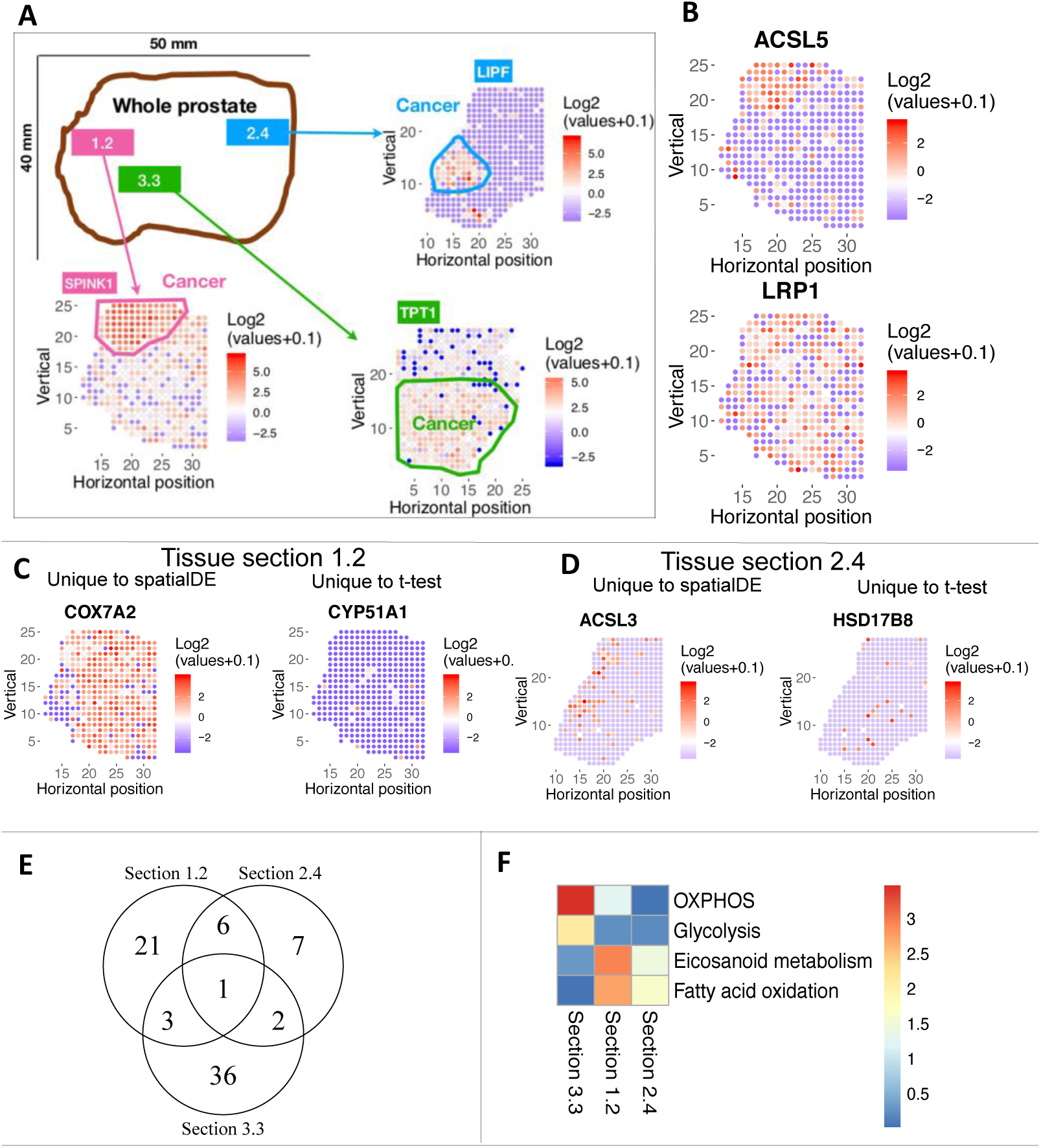
Spatially variable metabolic gene expression across three tissue sections of a prostate cancer patient. **A.** Overview of the previously published spatial transcriptomics dataset used in the study. The malignant region is circled in each tissue section. Each spot in a biopsy is 100 μm diameter; adjacent spots are 200 μm center-to-center. **B.** Example of a spatially variable gene, ACSL5; and a gene that is not spatially variable, LRP1. Each dot represents a different locus at which gene expression was profiled. The colors correspond to the log2 transformation of normalized expression values across each tissue section. Red color denotes higher expression; blue denotes lower expression. **C and D:** Compared to t-test, spatially variables genes found by spatialDE tend to have spatial continuity. C: examples in section 1.2, COX7A2 is only found by spatialDE, while CYP51A1 is only found by t–test. D: examples in section 2.4. ACSL3 is only found by spatialDE, while HSD17B8 is only found by t-test. **E.** Venn diagram of spatially variable genes in three tissue sections. Majority of spatially variable genes are specific to a tissue section. **F.** Metabolic pathway enrichment of spatially variable genes in each tissue section. Color indicates negative log10 of enrichment p-value. There are 64 metabolic pathways in total. The four pathways with significant enrichment in at least one tissue section are shown.

We also compared SV genes (i.e., those identified by spatialDE) to genes identified by t-test as differentially expressed between tumor vs. non-tumor regions (defined in Figure 1A). Genes uniquely discovered by spatialDE tend to have a spatially clustered structure (i.e., spatial continuity, left panels of Figure 1 C and D). On the other hand, differentially expressed genes uniquely found by t-test tend to lack spatial continuity and show scattered expression (right panels of Figure 1 C and D). Thus, we used spatialDE throughout the following analysis. It is worth noting, however, that many SV genes are not captured by t-test (Figure S1A) and that SV genes need not to be restricted to tumor vs. normal comparison *a priori*; e.g., COX7A2 is expressed in both tumor and prostate intraepithelial neoplasia (PIN) regions and depleted in normal prostate gland (Figure 1C and Figure 2A).

**Figure 2.**
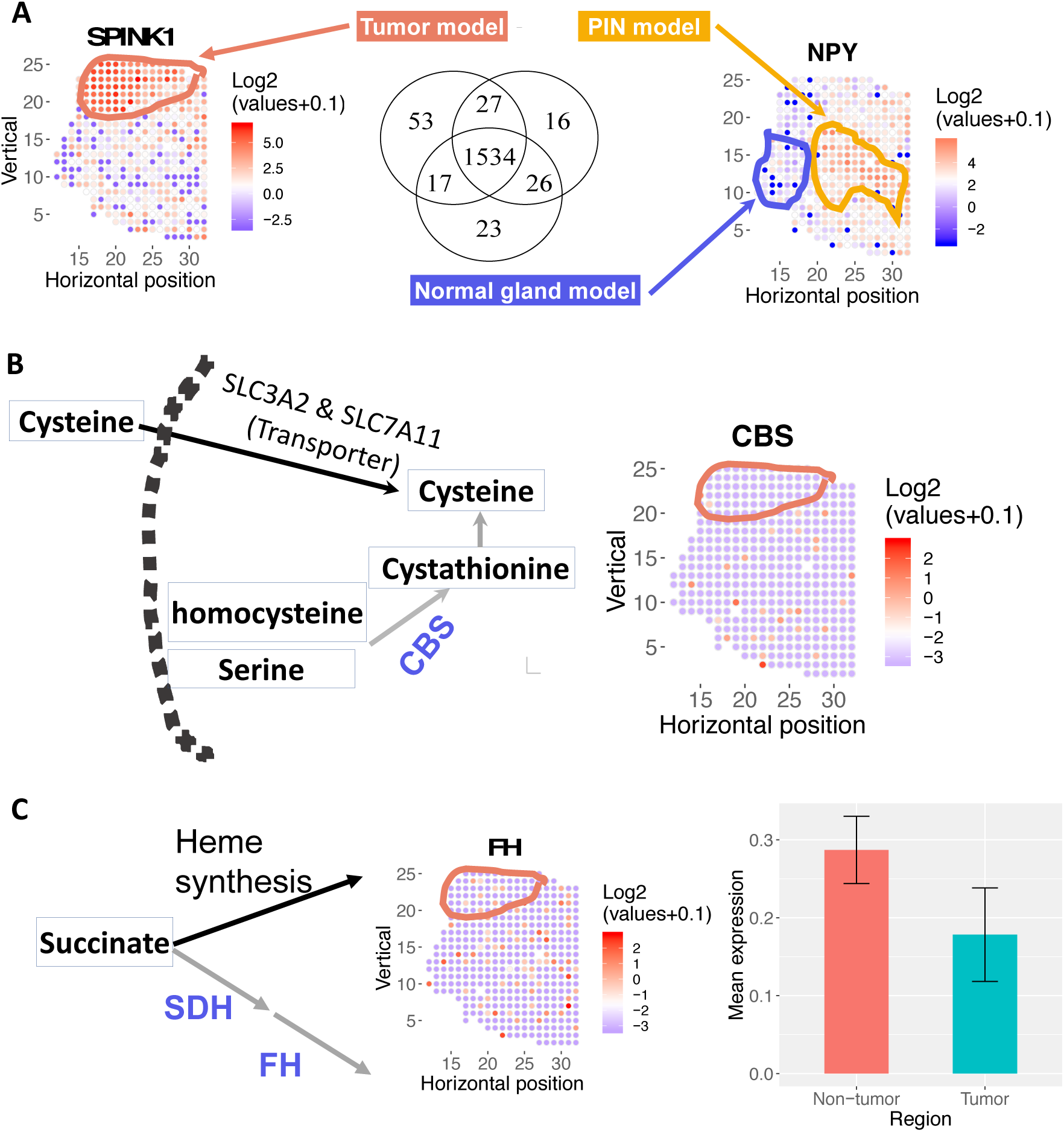
Spatially resolved-metabolic network models for tissue section 1.2 and mechanistic predictions. **A.** Number of overlapping reactions for each region-specific metabolic network model. Mean expression profiles across locations circled for each region are used to build the models, via the mCADRE algorithm. mCADRE extracts a tissue-specific sub-network from the input generic human metabolic network model based context-specific transcriptomic data^25^. Locations not outlined as tumor, normal or PIN (prostatic intraepithelial neoplasia) are enriched for stromal markers, based on^12^. SPINK1 and NPY gene expression mark malignant cells and PIN respectively. **B.** Model predicts that disrupting cysteine transport via SLC3A2 and SLC7A11 is selectively lethal in tumor region because *de novo* synthesis via the CBS gene is depleted in the tumor region. Left: metabolic pathway diagram. Each rectangle represents a metabolite. Each arrow represents a reaction or transport (black arrow: reaction is present in the tumor; gray arrow: reaction is absent from the tumor). The name of each reaction is labeled above the corresponding arrow, and CBS is highlighted in blue. The dashed arc represents the plasma membrane. Right: log2 transformation of normalized expression values of CBS across the tissue section. Red means higher expression; blue/white means low or no expression. **C.** Model predicts that disrupting succinate utilization via heme synthesis and degradation is lethal in tumor region because fumarate hydratase is depleted in the tumor region. Left: metabolic pathway diagram. Each rectangle represents a metabolite. Each arrow represents a reaction or transport (black arrow: reaction is expressed in tumor; grey arrow: reaction is absent in tumor), the name of each reaction is labeled above the corresponding arrow. Middle: log2 transformation of normalized expression values of FH across the tissue section. Right: Mean expression of FH in non-tumor and tumor region. Error bar represents standard error of the mean.

Interestingly, most SV genes are unique to each tissue section (Figure 1E), potentially because tumor cells from different regions of the prostate developed distinct survival strategies. Only one gene–Acid Phosphatase, Prostate (ACPP)–is spatially variable in all three tissue sections. ACPP is a known prostate cancer marker^20^, but spatial transcriptomics data suggest that ACPP is only enriched in the tumor region in section 3.3. It is enriched in *non-tumor* regions in section 1.2 and 2.4. (Figure S1B). This highlights the spatially heterogeneous expression pattern of this known marker gene that would have been missed by bulk averaging of the whole biopsy.

Metabolic pathway enrichment analysis also showed that SV genes are enriched in arachidonic (i.e., eicosanoid) and fatty acid metabolism in section 1.2, while SV genes are enriched in glycolysis and OXPHOS in section 3.3 (Figure 1F). Notably, the mean expression profiles of SV genes in glycolysis and OXPHOS are both high in the region surrounding the malignancy, and *low* in the malignant region itself (Figure S1C and D). This is consistent with previous findings that, unlike other cancer types, early stage primary prostate cancer is known to *not* exhibit elevated glucose consumption (i.e., does not exhibit the Warburg effect)^3^. Our analysis further showed that certain primary prostate cancer cells have lower glycolysis and OXPHOS activities than their adjacent normal tissues, and thus may not respond as effectively to glycolysis or OXPHOS inhibitors. However, as prostate cancer cells become more invasive and metastatic at later stages, the glycolysis pathway is up-regulated^3, 21^.

Our analysis also revealed interesting spatial patterns of reactive oxygen species (ROS) gene expression. SOD2 (superoxide dismutase 2), which protects mitochondria from reactive oxygen species, including those generated by OXPHOS complexes^22^, has a spatial pattern that is opposite of OXPHOS genes in section 3.3 (Figure S1D and E). This suggests that certain prostate cancer cells are under higher ROS stress despite lower OXPHOS activity, and thus require higher expression of SOD2. Therefore, targeting the ROS detoxification machinery may selectively kill these cancer cells. On the other hand, SOD3, which is an extracellular superoxide dismutase, has the same spatial distribution as OXPHOS expression (i.e., lower in tumor region, higher in adjacent non-tumor region, Figure S1D and E). This agrees with previous reports that loss of SOD3 expression has been shown to promote cancer cell migration and invasion, including in prostate cancer^23^, and increasing SOD3 expression has been shown to improves tumor response to chemotherapy by regulating endothelial cell structure and function^24^. Thus, spatially resolved transcriptomics data can be used to guide whether patients will respond to drugs that increase SOD3 levels (e.g., by the FDA-approved drug Lovastatin) and synergize with chemotherapy^24^.

Taken together, the data suggest that there is significant spatial heterogeneity of metabolic gene expression within the same tumor biopsy, and such spatial metabolic heterogeneity can be exploited to guide targeted therapy.

### Intra-biopsy tumor metabolic heterogeneity presents new selective metabolic targets in cysteine and succinate metabolism

To further elucidate the spatial patterns of metabolic gene expression, and identify opportunities to selectively target metabolic aberrations in malignant cells, we built spatially resolved metabolic network models of each tumor and no-tumor region in each tissue section using the mCADRE algorithm^25^. mCADRE infers a tissue-specific metabolic sub-network based on context-specific transcriptomic data and the input generic human metabolic network. It has been independently validated as achieving high accuracy predicting lethal cancer metabolic genes^26^.

After identifying what metabolic genes are spatially variable, we further identified *where* these SV metabolic genes are expressed. Cancer, prostate intraepithelial neoplasia (PIN), and normal prostate gland regions are outlined based on computational inference and IHC staining in Berglund *et al* ^12^. We built a genome-scale metabolic network model (GEM) for each region (Figure 2A), and systematically simulated how knocking down each metabolic gene affects proliferation. We identified 16 genes whose *in silico* knockdowns are selectively lethal for malignant cells using the tumor-specific model but are missed by a model built using the mean transcriptome of all spatial locations within the tissue section (i.e., “pseudo-bulk data”, Table 1). Malignant-, normal-specific and “pseudo-bulk” GEMs provide potential mechanistic explanations for why these genes may be selectively lethal in malignant cells. Figure 2B and 2C provide two such examples discussed below.

**Table 1.** 16 metabolic genes predicted to be selectively lethal based on tumor-specific model but missed by the pseudo-bulk model for prostate cancer tissue section 1.2. Column 1 lists genes specifically lethal in the tumor-specific model but not the pseudo-bulk model built with mean expression across all spatial locations. Column 2 lists the pathways of genes in column 1. Column 3 lists genes that are not present in the tumor-specific model but present in the bulk model and underlie the differential lethality of genes in column 1 (i.e., genes in column 3 provide an alternative metabolic path).

| **Genes lethal in tumor not in bulk model** | **Pathways** | **Alternative gene not in tumor model** |
| --- | --- | --- |
| CPOX | Heme synthesis | FH |
| HMBS | Heme synthesis |  |
| ABCC1 | Heme synthesis |  |
| PPOX | Heme synthesis |  |
| UGDH | Heme synthesis |  |
| UROD | Heme synthesis |  |
| UROS | Heme synthesis |  |
| SLC35D1 | Heme synthesis |  |
| UGT1A8 | Heme synthesis |  |
| SLC3A2 | Cysteine metabolism | CBS |
| SLC7A11 | Cysteine metabolism |  |
| AACS | Acetyl-Acetate-CoA | ACAT2 |
| SLC6A14 | Tryptophan transport | SLC16A10 |
| SLC6A12 | Glycine betaine transport | BHMT |
| FASN | De novo fatty acid synthesis | ADK1 |
| SLC25A1 | De novo fatty acid synthesis |  |

#### Cysteine

Our metabolic model simulations predict that malignant cells are selectively vulnerable to the knockdown of the cysteine transporter complex that consists of the transporters SLC3A2 and SLC7A11, because the enzyme for cysteine *de novo* biosynthesis, cystathionine-beta-synthase (CBS), is selectively depleted in malignant cells (Figure 2B, CBS expression; Figure S2A, bar plot of CBS-expressing, i.e., expression level >0, locations in tumor and non-tumor regions). Cysteine depletion by inhibiting the cysteine-glutamate antiporter xCT using Sulfasalazine (SSZ) has been previously shown to markedly inhibit the proliferation of prostate cancer cell lines DU-145 and PC-3 *in vitro*^27^. Our models suggest that cysteine depletion can also selectively affect malignant cell growth *in vivo* due to loss of *de novo* synthesis. Since SSZ is already approved by the FDA to treat rheumatoid arthritis, ulcerative colitis, and Crohn’s disease, it is attractive to re-purpose it for prostate cancer treatment. SSZ may be more effective in androgen-independent prostate cancer cells where CBS expression is lower^28^.

#### Succinate

Succinate is a key intermediate in the TCA cycle. Our model predicts that malignant cells are selectively vulnerable to the inhibition of the heme synthesis pathway because fumarate hydratase and succinate dehydrogenase are selectively depleted in malignant cells (Figure 2C and Figure S2B). Fumarate hydratase and succinate dehydrogenase are known tumor suppressors^29^. It has been previously shown that inhibiting heme synthesis is selectively lethal to renal clear cell carcinoma with fumarate hydratase mutation^17^. Our model suggests that this synthetic lethal interaction may also be exploited in prostate cancer. This is especially interesting given that somatic mutations in fumarate hydratase have been reported in a small subset of prostate cancer patients^30^. Succinate metabolism is also spatially variable in tissue section 2.4. Our model also predicts that *GTP-specific* beta subunit of succinyl-CoA synthetase (SUCLG2) is selectively lethal in malignant prostate cancer cells because the alternative route via *ATP-specific* succinyl-CoA synthetase (SUCLA2) is absent in the malignant model (Figure S2C). SUCLA2 has been previously reported to be significantly down-regulated in prostate cancer^31^. Our model predicts that SUCLA2 down-regulation creates a selective vulnerability to SUCLG2 knockdown in malignant cells.

These results demonstrated that metabolic network models based on spatial transcriptomics data can reveal novel selective metabolic vulnerabilities that are missed by models based on bulk gene expression data from entire tissue biopsies.

### Spatial heterogeneity of fatty acid metabolism in the tumor microenvironment presents new selective targets

Spatially variable genes in tissue section 1.2 are enriched for fatty acid (FA) and arachidonic acid metabolism (Figure 1F). Furthermore, our tumor-specific model also predicts that perturbations in multiple genes of the fatty acid synthesis pathway are selectively lethal in malignant cells (Table 1). Given that dysregulation of lipid and fatty acid metabolism is a major feature of prostate cancer^4, 32^, we further explore spatial heterogeneity of FA and lipid metabolism using spatially-resolved metabolic network models.

#### Cholesterol synthesis

Acetoacetate-CoA is a precursor for cholesterol synthesis, an essential component of cellular membranes. Our model predicts that Acetoacetyl-CoA Synthetase (AACS) depletion is selectively lethal to malignant cells because the alternative route for Acetoacetate-CoA synthesis, Acetyl-CoA Acetyltransferase 2 (ACAT2) is selectively depleted in the tumor region (Figure 3A). ACAT2 is known to be down-regulated in prostate cancer^33^. This selective prediction is missed by the pseudo-bulk model.

**Figure 3.**
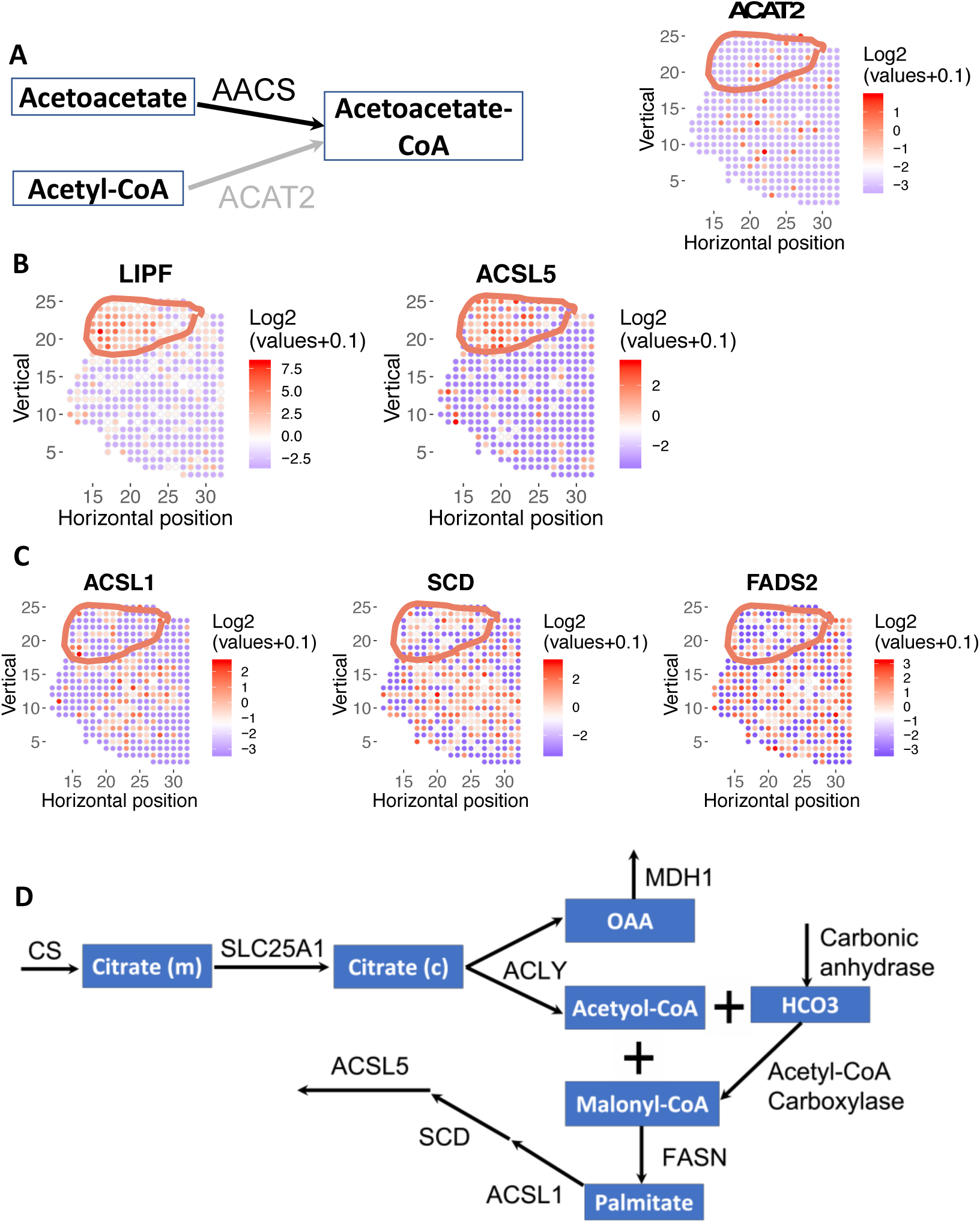
Fatty metabolism is spatially variable in prostate cancer. Tumor region is circled. **A.** Model predicted that AACS is lethal in tumor because the alternative route for acetoacetate-CoA synthesis, ACAT2, is depleted in tumor region. Left: metabolic pathway diagram. Each rectangle represents a metabolite. Each arrow represents a reaction or transport (black arrow: reaction is present in the tumor; gray arrow: reaction is absent from the tumor), the name of each reaction is labeled above the corresponding arrow. Right: log2 transformation of normalized expression values across the tissue section. Red means higher expression; blue/white means low or no expression. **B.** Lipid hydrolysis (via Lipase F) and fatty acid synthesis (via ACSL5) is highly enriched in the tumor region. **C.** Fatty acid oxidation (via ACSL1) and desaturation (via SCD and FADS2) are depleted in tumor region and higher in PIN and normal region. Color denotes log2 transformation of normalized expression values of a gene of interest. **D.** A schematic of all genes and reactions predicted to be essential for metabolic flux through the reaction catalyzed by the tumor-enriched gene ACSL5. Each rectangle represents a metabolite. Each arrow represents a reaction or transport, the name of each gene/reaction is labeled above the corresponding arrow (gene names: CS, SLC25A1, ACLY, MDH1, FASN, ACSL1, SCD, ACSL4; reaction names: carbonic anhydrase, acetyol-CoA carboxylase).

#### Fatty acid metabolism

We found that hypoxia potentially explains the spatially distinct distributions of fatty acid metabolic genes. Metabolic genes in lipolysis (LIPF), and fatty acid synthesis (ACSL5) are selectively expressed in the malignant region (Figure 3B and S3A & B). Recent studies showed that prostate cancer cells show elevated uptake of extracellular fatty acids^4, 32^. Our analyses suggest that free fatty acids generated by lipolysis via LIPF can be a potential source of extracellular free fatty acids. In contrast to tumor-enriched lipolysis and fatty acid synthesis enzymes that do not require oxygen, fatty acid metabolic genes that require molecular oxygen are depleted in the tumor region, including fatty acid desaturation (SCD, FADS2) and oxidation (ACSL1) (Figure 3C & S3C). Although ACSL1 and ACSL5 are isozymes with similar catalytic function, genetic knockout studies in mice showed that ACSL5 has a major role in fatty acid biosynthesis and deposition, while ACSL1’s function is mostly involved in fatty acid oxidation^34^. A metabolic model based on bulk gene expression data would incorrectly assume that both enzymes are expressed by malignant cells, thus over-estimating the metabolic capabilities of malignant cells.

We identified additional selective metabolic liabilities that are driven by malignant cells’ dependence on *de novo* fatty acid synthesis, by maximizing metabolic flux through the tumor-enriched ACSL5 reaction in our model (Figure 3D). Reassuringly, we recovered several genes involved in *de novo* fatty acid synthesis, specifically citrate synthase (CS), mitochondrial citrate transporter (SLC25A1), ATP citrate lyase (ACLY), and fatty acid synthase (FASN). We also identified additional selective liabilities, specifically, ACSL1, cytosolic malic dehydrogenase (MDH1), carbonic anhydrase, and stearoyl CoA desaturase (SCD). ACSL1 has been previously shown to be important for biosynthesis of C16:0−, C18:0−, C18:1− and C18:2-CoA, triglycerides and lipid in prostate cancer cells and ACSL1 knockdown inhibited prostate cancer cell proliferation and migration *in vitro* and *in vivo*^35^. In addition to knockdown, ACSL1 can also be pharmacologically inhibited by small molecules Triacsin C^36^. Carbonic anhydrase has been previously reported to be important for de novo lipogenesis^37^. SCD1 produces monounsaturated fatty acids from saturated fatty acids, and has been shown to be important for cancer initiation, proliferation, and metastasis in many types of cancer, including prostate cancer, and can be inhibited using small molecules such as CAY10566 and TOFA^38–40^. Thus, the role of ACSL1, carbonic anhydrase, and SCD1 in cancer are all supported by literature. Although MDH1 inactivation inhibits pancreatic cancer growth by suppressing glutamine metabolism^41^, the role of MDH1 in *de novo* fatty acid synthesis has not been previously studied, and may be a potential new target to manipulate fatty acid metabolism for prostate cancer treatment.

### Spatial patterns of arachidonic acid metabolism

Arachidonic acid is the starting point for the synthesis of prostaglandins and leukotrienes, both of which have immunomodulatory functions^42^. We identified enzymes in arachidonic acid metabolism that show spatially distinct expression patterns, as well as selective targets to disrupt prostaglandin synthesis from arachidonic acids (Figure 4). PTGDS (Prostaglandin D2 Synthase) and HPGD (15-Hydroxyprostaglandin Dehydrogenase) are enriched in the tumor region, while MGST3 (Microsomal Glutathione S-Transferase 3) is depleted in the tumor region in tissue section 1.2 (Figure 4A and S4A). The reaction network formed by these enzymes is depicted in Figure 4A. HPGD, MGST3 and other arachidonic acid metabolic genes also show spatially distinct expression patterns in other tissue sections (Figure S4B and C).

**Figure 4.**
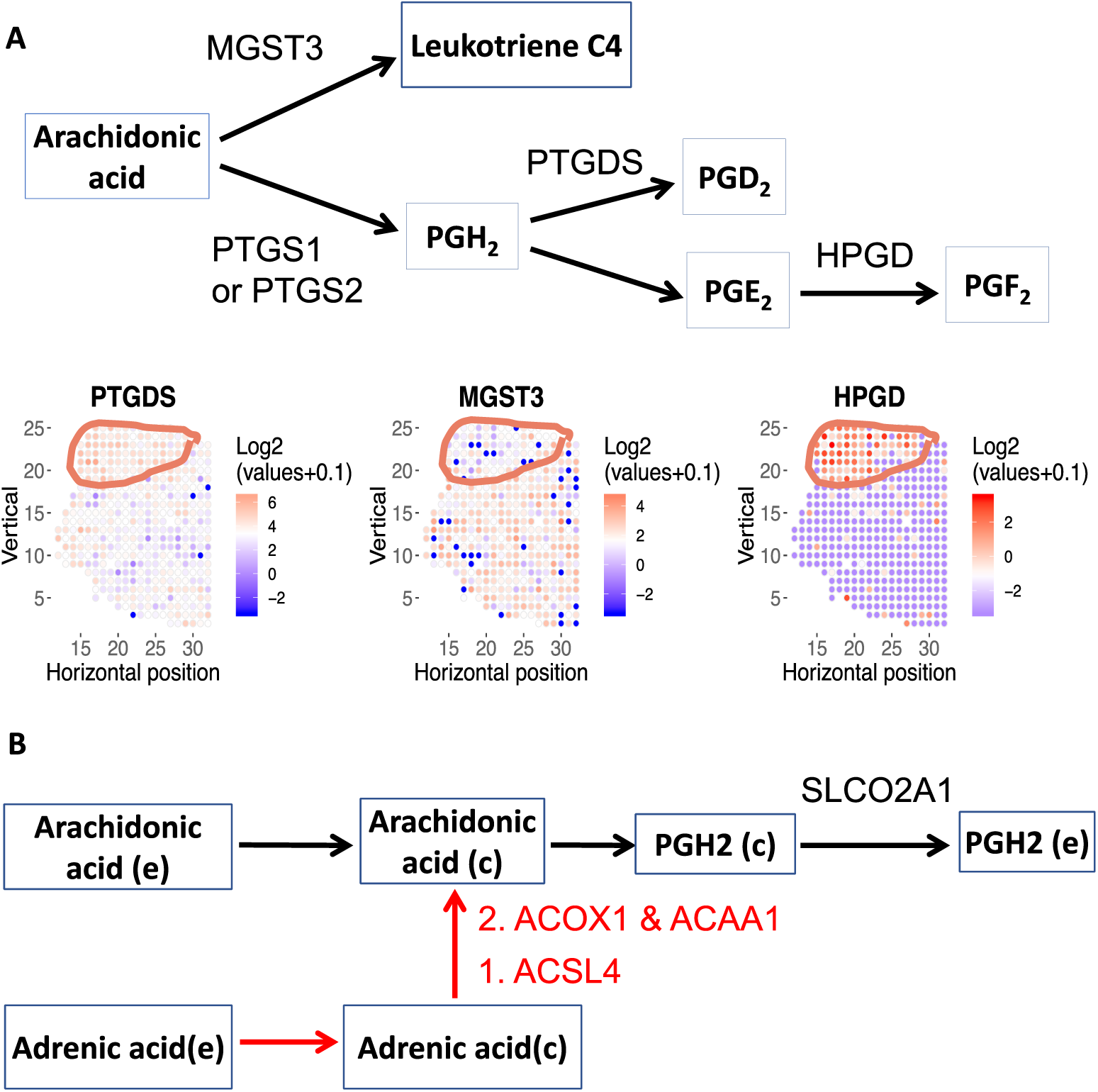
Arachidonic acid metabolism is spatially variable. **A.** Metabolic genes in prostaglandin and leukotriene synthesis are spatially variable in tissue section 1.2. While PTGDS and HPGD are enriched in tumor region, MGST3 is selectively depleted in tumor region. HPGD is also spatially variable in tissue section 2.4; PTGS2 (i.e., COX-2) is spatially variable in tissue section 3.3 (Figure S4B). Top: metabolic pathway diagram. Each rectangle represents a metabolite. Each arrow represents a reaction or transport, the name of each reaction is labeled above the corresponding arrow. Bottom: Expression level of 3 arachidonic acid metabolism genes. Red color denotes higher expression; blue denotes lower expression. **B.** Model predicted that SLCO2A1 is essential for arachidonic acid metabolism. In addition to arachidonic acid, adrenic acid can also contribute to prostaglandin metabolism. Each rectangle represents a metabolite. Each arrow represents a reaction or transport, the name of each reaction is labeled above the corresponding arrow. Genes involved in conversion of adrenic acid to arachidonic acid is highlighted in red.

Previous analyses have shown that arachidonic metabolism is dysregulated in multiple types of cancer^25, 43^, and inhibition of key arachidonic metabolic genes results in massive apoptosis in prostate cancer cells^44^. The distinct spatial expression patterns of arachidonic acid metabolism genes imply that different molecular species of prostaglandin and leukotrienes are enriched or depleted in the malignant region. MGST3 is used for the synthesis of leukotriene C4, a major mediator of endoplasmic reticulum stress and oxidative DNA damage^45^. Our analysis suggests that leukotriene C4 is depleted in malignant cells. HPGD catabolizes prostaglandin E2 (PGE^2^) into PGF^2^. Intriguingly, while HPGD has been widely reported as a tumor suppressor in multiple types of cancer^46–48^, it is selectively enriched in the malignant cells of both tissue sections 1.2 and 2.4. HPGD expression is induced by androgen and is up-regulated in the androgen-dependent prostate cancer cell line LNCaP ^49^. Because PGE^2^ has angiogenic^50^ and immunosuppressive functions^51^, higher HPGD expression indicates that the malignant region is depleted of PGE^2^ and more amenable to cancer immunotherapy.

Since the reaction catalyzed by PTGS1 and 2 (Prostaglandin-Endoperoxide Synthase 1 and 2, commonly known as COX-1 and COX-2) is the first step in prostaglandin synthesis and known to be up-regulated in prostate cancer^52^, we used our tumor-specific metabolic network model to simulate additional metabolic liabilities that are driven by the PTGS reaction (Figure 4B). We found that SLCO2A1 is essential for the PTGS reaction. Blocking SLCO2A1 has been shown to reduce colon cancer tumorigenesis^53^. Importantly, SLCO2A1 can be potently and selectively inhibited by the FDA approved drug suramin^54^. Therefore, SLCO2A1 may be an attractive target in prostate cancer. Arachidonic acid is required for prostaglandin synthesis. In addition to arachidonic acid uptake, our model simulation also revealed that cancer cells can use adrenic acid as an alternative source of arachidonic acid. Our model predicted that adrenic acid can be converted to arachidonic acid via reactions catalyzed by ACSL4, ACOX1, and ACAA1 (Figure 4B). In particular, ACOX1 is selectively up-regulated in HER2-positive subtypes of breast cancer and is positively associated with shorter survival^55^ and may be a potential target in prostate cancer. In addition to cancer-intrinsic functions, arachidonic acid uptake and synthesis of prostaglandins such as PGE^2^ have immunosuppressive functions^51^. Because both adrenic acid and arachidonic acid are present in prostate cancer specimens^56^, the adrenic-to-arachidonic pathway suggests that blocking both arachidonic *and* adrenic uptake may be required to abolish the immunosuppressive effects of PGE^2^.

### Spatial patterns of arginine and urea metabolism

Arginine metabolism is dysregulated in a wide range of cancers, and arginase is an attractive drug target^57^. One product of arginase is urea, and we found that the urea transporter SLC14A1 is selectively depleted in the malignant region in both tissue sections 1.2 and 2.4 (Figure 5A). More importantly, we also found that SLC14A1 is significantly lower in PIN, prostate cancer *in situ* and metastatic prostate cancer compared to normal prostate by re-analyzing a large set of prostate cancer patients^58^ (Figure 5B). SLC14A1 is also down-regulated in lung, prostate and urothelial cancer^59^. Arginine catabolism by arginase generates ornithine, a key substrate for polyamine synthesis, which has important signaling functions in prostate cancer^60^. Alternatively, arginine is also important for biosynthesis, which creates a competition for polyamine synthesis (Figure 5C). Through model simulation, we found that increased flux through the urea transport reaction leads to decreased growth (Figure 5D). Thus, down-regulation of urea transport is a strategy by malignant cells to use arginine for growth. Therefore, inducing SLC14A1 expression is a potential strategy to inhibit prostate cancer growth. Indeed, transfection of SLC14A1 into lung cancer cell line H520 inhibited colony formation^61^. Our model simulation also found that, to compensate for increased flux via SLC14A1 and maintain growth rate, cells need to increase the uptake of arginine by 4- to 5-fold (Figure 5E). Thus, our model predicts that induction of SLC14A1 expression, combined with arginine depletion may kill cancer cells.

**Figure 5.**
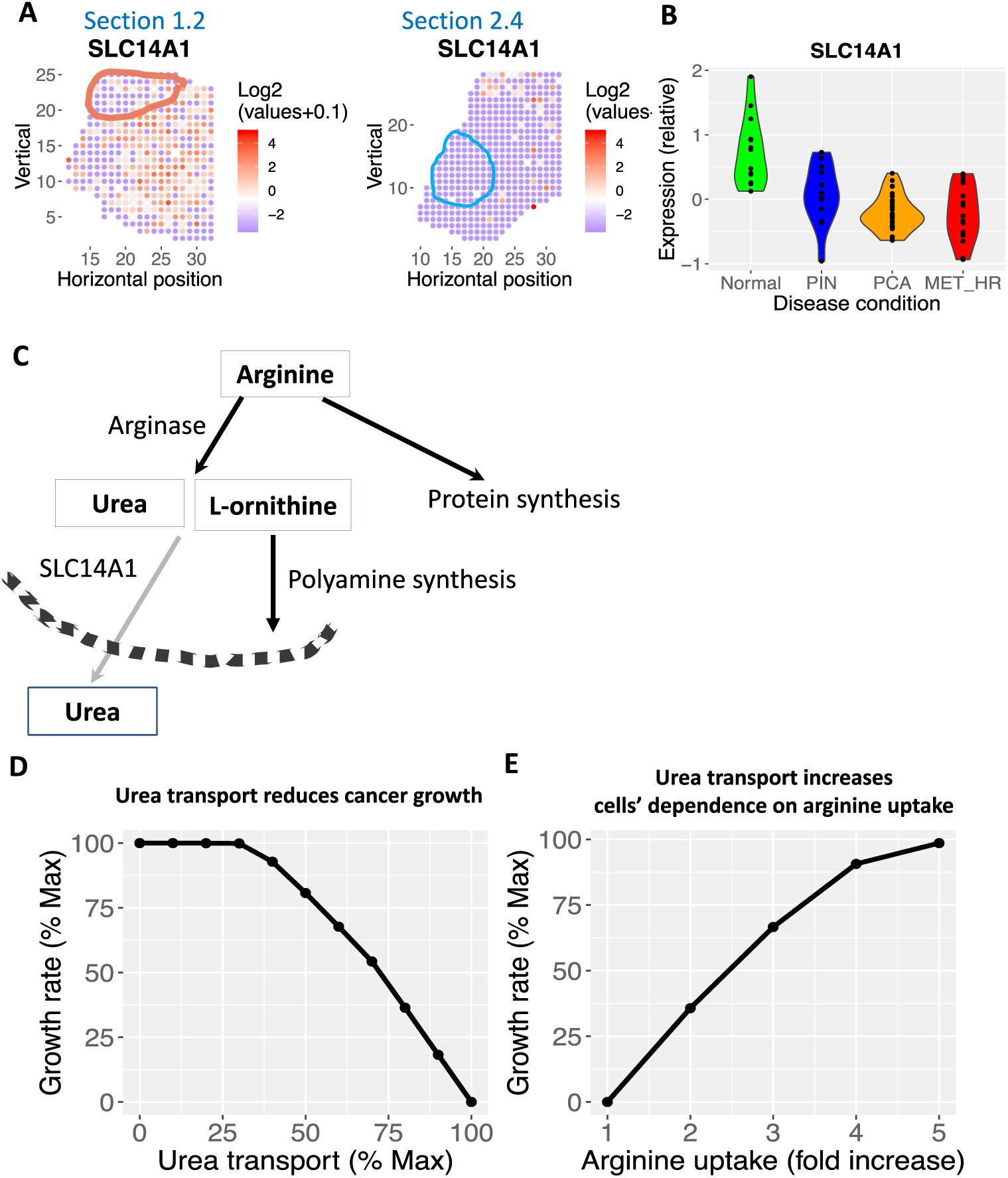
Urea transport is spatially variable in prostate cancer. **A.** The urea transporter, SLC14A1, is depleted in tumor region and highly expressed in non-malignant region in both tissue section 1.2 and 2.4. Red color denotes higher expression; blue denotes lower expression. Tumor regions are outlined. **B.** SLC14A1 is also down-regulated in another prostate cancer study using laser-capture micro-dissected normal gland, PIN region, prostate cancer (PCA) and metastatic prostate cancer (MET-HR). **C.** Arginine can be used for growth or arginase reaction. While arginase reactions produce an essential substrate of polyamine synthesis (L-ornithine), the other product, urea, are transported out of cells by SLC14A1. Each rectangle represents a metabolite. Each arrow represents a reaction or transport, the name of each reaction is labeled above the corresponding arrow. The dashed arc represents the plasma membrane. **D.** Model predicted that higher flux through urea transport is correlated with reduced growth rate. x-axis, flux through the urea transport reaction as a percentage of maximum feasible flux; y-axis: growth rate as a percentage of maximal growth rate. **E.** Model predicted that, at maximum urea transport flux, cells need to increase arginine uptake flux by 4-5 fold in order to restore growth rate. x-axis, flux through the arginine uptake reaction as a percentage of maximum feasible flux; y-axis: growth rate as a percentage of maximal growth rate.

## Discussion

The genome-scale metabolic network models of prostate cancer developed by our novel pipeline using spatially-resolved transcriptomics data have revealed many new, malignancy-specific metabolic perturbations that would have been missed by models based on bulk gene expression data of the whole tissue biopsy. Our model predictions span amino acid (cysteine and arginine/urea), fatty acid and lipid (cholesterol, fatty acid synthesis/oxidation, arachidonic acid) metabolism, and the TCA cycle (succinate). Many of our predictions are supported by previous literature, which provides further confidence to explore the novel predictions as potential drug targets for prostate cancer. Importantly, many of the metabolic genes predicted to be selectively lethal in prostate cancer cells can be targeted by FDA-approved small molecule compounds.

Unlike other solid tumors, primary prostate cancer does not exhibit the classical Warburg effect (i.e., does not exhibit elevated glycolysis). Instead, prostate cancer shows elevated *de novo* fatty acid and lipid synthesis^62^. Recent evidence also demonstrates that extracellular fatty acids are major contributors to lipid synthesis in prostate cancer^32^. In fact, prostate cancer cells show higher uptake of fatty acids than glucose, especially in metastatic and circulating prostate cancer cells^63, 64^. Suppressing fatty acid uptake via CD36 has also been shown to inhibit prostate cancer growth^4^. However, the sources of free fatty acids are not fully characterized. Our modeling analysis suggests that Lipase F (LIPF) can potentially degrade extracellular triglycerides and generate free fatty acids for cancer cells to uptake (Figure 3B). Thus, targeting extracellular lipid degradation may inhibit prostate cancer growth.

In addition to the uptake of extracellular free fatty acid through LIPF and CD36, our model also suggested that prostate cancer cells also exhibit elevated *de novo* fatty acid synthesis. Our analysis finds that ACSL5 gene is strongly enriched in one tumor region (section 1.2), and it has been also been found to be over-expressed in other prostate cancer^65^. ACSL5 plays a critical role in lipid droplet formation ^66^, and lipid droplet formation promotes prostate cancer aggressiveness^67, 68^. Therefore, targeting ACLS5 may be a potential strategy to inhibit the formation of lipid droplet formation and prostate cancer cell survival.

Hypoxia is a prominent feature of the tumor microenvironment, and malignant cells adapt their metabolic profiles to survive in the hypoxic environment^69^. Fatty acid desaturation, which requires molecular oxygen, is inhibited in hypoxic tumor regions (SCD and FADS2, Figure 3B). SCD is the best-known route to fatty acid desaturation. A recent study found that cancer cells can bypass SCD by using FADS2 for fatty acid desaturation^70^. However, our analysis of the spatial transcriptomics data revealed that both SCD and FADS2 are depleted in the malignant region. Inactivation of fatty acid desaturation creates the need to uptake exogeneous unsaturated fatty acids in order to maintain correct composition of saturated vs. unsaturated lipids in biological membranes^69,71^. Although exogeneous fatty acid uptake has been shown to be important for prostate cancer^4, 32^, the relative importance of exogeneous saturated vs. unsaturated fatty acids has not been examined^32^. Our model predicts that malignant cells will be more sensitive to depletion of exogeneous *unsaturated* fatty acids due to defective endogenous desaturation. Therefore, inhibition of exogeneous fatty acid uptake by targeting CD36 may synergize with inhibition of desaturation by targeting SCD (e.g., via small molecules CAY-10566 and TOFA^39, 40^) in killing hypoxic cancer cells. Hypoxia also induces lipid droplet formation by up-regulating the expression of fatty acid transporters via HIF-1a^72^.

Besides fatty acid desaturation and lipid droplet formation, our analysis also suggests that the fluctuating oxygen levels in the TME^73^ could also sensitize prostate cancer cells to the inhibition of the mitochondrial citrate transporter SLC25A1 (Figure 3D). Our model’s prediction of SLC25A1’s essentiality in hypoxic tumor cells is substantiated by prior findings that SLC25A1 expression is up-regulated when prostate cancer cells are exposed to cycling hypoxia/re-oxygenation stress^74^ Notably, pharmacological inhibition of SLC25A1 sensitizes cancer cells to ionizing radiation, cisplatin or EGFR inhibitor treatments in lung cancer^74, 75^. Thus, treating prostate cancer cells in a hypoxic TME with the SLC25A1 inhibitor, 1,2,3-benzene-tricarboxylic acid (BTA) could not only yield direct tumor-specific killing, but it could also potentiate the activity of concomitant radiation or chemotherapy interventions.

Arachidonic acid is a potent signaling lipid, and precursor to the synthesis of a wide range of other signaling lipids such as prostaglandins and leukotrienes. Prostaglandin and leukotriene C4 have important functions in angiogenesis and immunomodulation. Our analysis showed that malignant cells have elevated synthesis of prostaglandins and decreased synthesis of leukotriene C4 (Figure 4A), which may influence the sensitivity to immunotherapy. Elevated prostaglandin synthesis may be targeted by suramin, a FDA-approved drug that potently and inhibits the prostaglandin transporter SLCO2A1^54^.

Cancer cells showed elevated dependence on multiple amino acids, including cysteine (Figure 2B), glutamine, aspartate, asparagine and arginine^76–78^. Arginase breaks down arginine into urea and ornithine. Arginase is an important regulator of the immune system^79^. Arginine deprivation via arginase activation suppresses anti-tumor T cell activity, so blocking arginase activity may improve tumor immunotherapy^80, 81^. We showed that the urea transporter SLC14A1 is selectively depleted in the malignant region, which is supported by additional transcriptomics data (Figure 5A & B). While decreased flux through urea transport is beneficial for biomass synthesis (Figure 5 C & D), the accumulation of urea may be toxic to cells^59^.

The spatial locations of cells have profound impacts on their function. In addition to prostate cancer, our modeling approach can be used to study spatial heterogeneity and coordination of metabolic activities in a wide range diseases where spatially resolved transcriptomic datasets are currently available, such as breast cancer^9^, pancreatic cancer^14^, melanoma^13^, and amyotrophic lateral sclerosis^82^. It can also be used to study spatial regulation of normal organ physiology in the liver^83^, heart^84, 85^ and kidney^86^.

This study has several limitations that need to be addressed in the future. First, the models are based only on transcriptomics data, which does not directly reflect metabolic activities. Thus, although we recapitulated known metabolic features of prostate cancer, other metabolic dysregulation may be missed. This can be addressed in the future as spatially-resolved proteomic^87^ and metabolomic^88^ technologies improve and such data can be incorporated into the metabolic network model. Indeed, new technologies already enables simultaneous measurement of spatially-resolved transcriptome and proteome^89^. Second, due to the relatively small number of expressed genes per location (around 3000 genes), we could not build a model for each individual location. Instead, we built separate, discrete genome-scale metabolic models for each region (tumor, normal, and PIN) that aggregate transcriptomics data over the spatial locations that span each region. Given that the expression profiles of many metabolic genes are likely to exhibit gradient-like patterns that change with the distance to the tumor core and/or vasculature, we anticipate that models based on our region-specific data aggregation represent useful approximations of the regions’ underlying metabolic states. In the future, we will develop a hierarchical approach that adaptively balances spatial resolution with the amount of data to better model metabolic spatial continuity. Lastly, this study is based on detailed spatial analysis of three biopsies from one prostate cancer patient. Given the highly heterogeneous nature of prostate cancer, analysis of a single individual does not represent a comprehensive survey of important selective metabolic target genes. However, many of the metabolic genes we identified as selectively lethal based on the prostate cancer spatial transcriptome do have strong functional support from literature (i.e., inhibition or induction known to affect cancer proliferation, Table 2). This suggests that the additional novel targets we identified may also be involved in prostate cancer. Moreover, many metabolic genes we identified as having spatially selective expression pattern also agree with much larger cohorts of laser-capture micro-dissected transcriptomes of prostate cancer patients (ACSL5^65^ in Figure 3, SLC14A1^58^ in Figure 5, and SUCLA2^31^ in Figure S2C). As spatially-resolved transcriptomics technology become even more powerful and accessible to more researchers, we anticipate that the computational workflow we developed will be applied on larger cohorts in the future.

**Table 2.**
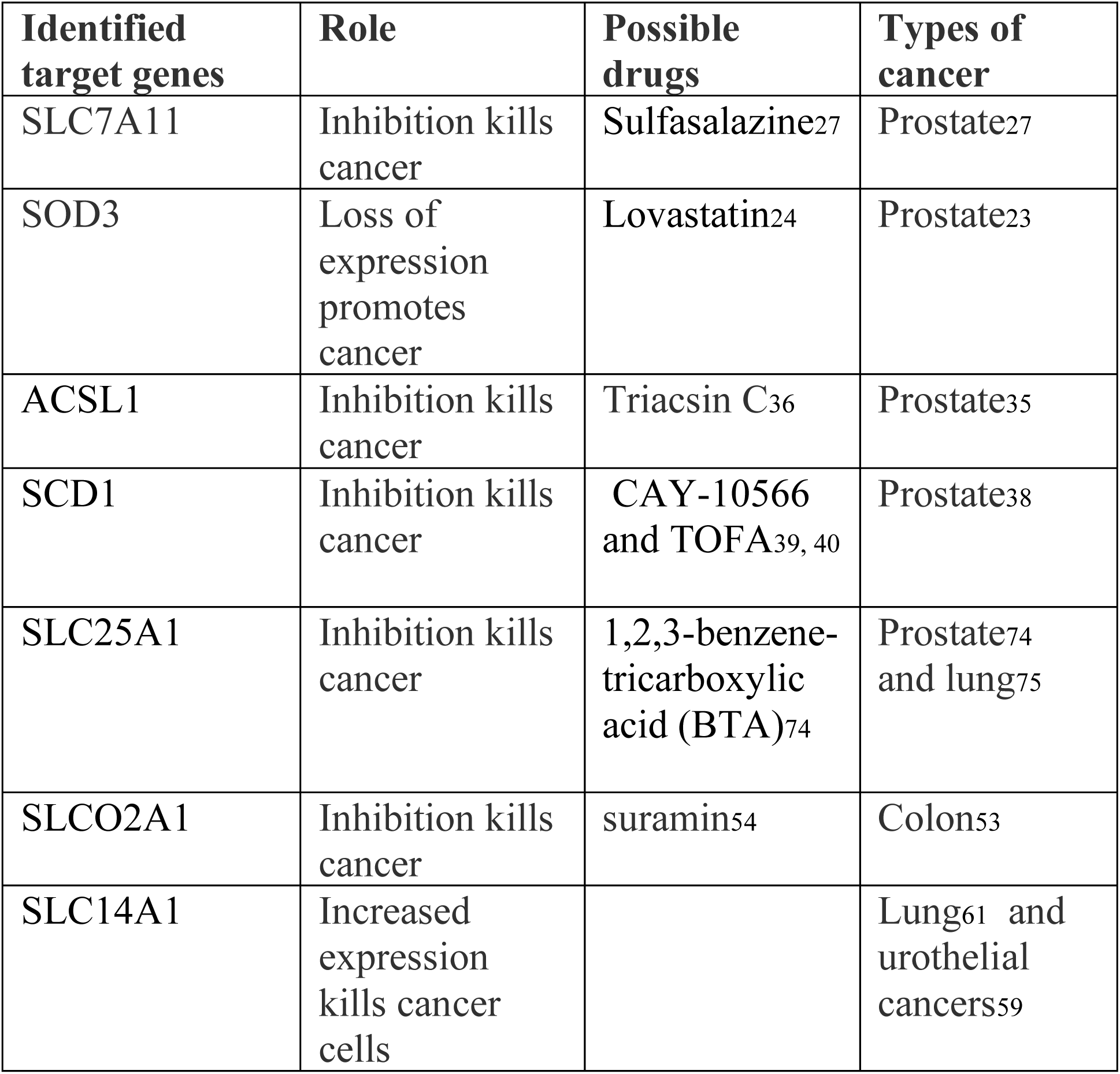
List of cancer-selective metabolic target genes identified by spatial metabolic analysis with literature support for their functional role in prostate or other types of cancer, as well as known inhibitors of these genes.

## Methods

The spatial transcriptomics dataset for prostate cancer^12^ was downloaded from the Spatial Transcritpomics Research website [http://www.spatialtranscriptomicsresearch.org/datasets/]. To identify spatially variable metabolic genes, we first extracted metabolic genes based on the latest version of the human metabolic network, Recon3D^90^. We used spatialDE^19^ to identify spatially variable genes. Briefly, we normalized expression data to the total read counts of all genes, removed metabolic genes with low expression, and we used spatialDE to find genes whose expression level at two locations depended on the distance between these two locations. spatialDE classifies genes into SV or non-SV by fitting two models: one where a gene’s expression covariance depended on location, and one without a spatial covariance matrix. If the former model fits better, then a gene is SV.

We used mCADRE^25^ to build genome-scale metabolic networks for normal, PIN and tumor regions. mCADRE has been previously validated as having good performance in predicting lethal metabolic genes in cancer^26^. mCADRE infers a tissue-specific metabolic network using context-specific transcriptomic data and a generic human metabolic network model. Some reactions involving multiple enzymes, such as enzyme complex formation, require the presence of all constituents, and are limited by the least abundant. By default, mCADRE modeled these by taking the min of the expression levels of the constituent genes. Given the sparsity of data (3000 detected genes per spatial location), taking the min for genes connected by AND (enzyme complexes) will result in mostly zeros. Therefore, we modified mCADRE to take the mean instead. Gene-level score was the mean expression level of a gene across all locations, not how often it is expressed above 0. Dually, we sum expression levels for genes connected by OR. A metabolic reaction is defined as a core reaction if it is expressed in 30% of all locations within each region. Remaining reactions are first ranked by their expression frequency across locations, then by their connectivity-based evidence. We removed highly connected metabolites such as H_2_O, ATP, ADP, Pi, NAD, NADH, etc., before calculating reaction connectivity. The COBRA Toolbox^91^ was used for gene knock out simulations.

### Model improvements

Recon 1^92^ (and Recon 2^93^, 3D^90^) all assumed SLC27A5 is the only transporter for arachidonic acid uptake. Latest evidence also demonstrates that SLC27A2 (FATP2) has a major influence for arachidonic acid uptake^51^. Latest evidence also suggests that ACSL4 (not ACSL1) favors arachidonic acid and adrenic acid as substrate^94^. We modified the gene-reaction rules to reflect both findings.

## Data availability statement

Prostate cancer spatial transcriptomics data used in this study can be found at: http://www.spatialtranscriptomicsresearch.org/datasets/10-1038-s41467-018-04724-5/ Bulk transcriptomics of prostate cancer samples are from Gene Expression Omnibus (accession number GSE6099).

R and Matlab codes, as well as spatially resolved metabolic networks for this project are deposited at Github (https://github.com/yuliangwang/prostate_cancer_spatial_metabolic_network.git)

## Acknowledgements

This project was funded by the University of Washington Royalty Research Fund to YW. We thank Peiyun Zhou for critical feedbacks on figures.

## Author Contributions Statement

Y.W. conceived the study and carried out the analyses. Y.W., S.M., and W.L.R. interpreted the results. Y.W. wrote the manuscript with input from S.M. and W.L.R.

## Competing Interests

The authors declare no competing interests.

## Supplemental figures

**Figure S1.**
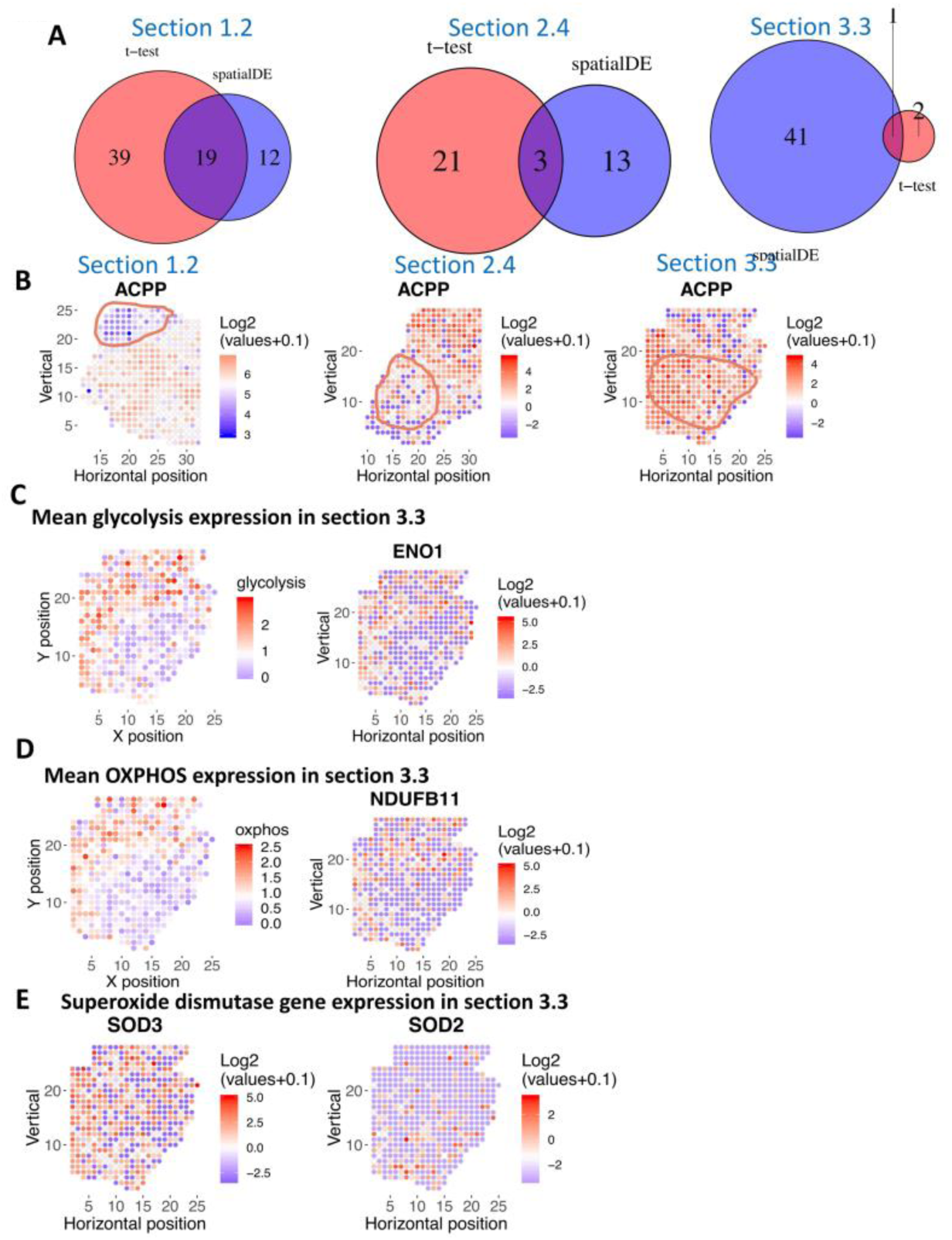
**A.** Overlap of genes identified as spatially variable by spatial DE vs. t-test. The overlap is bigger when tumor region is well-defined and clustered together. **B.** Log2 expression level of the ACPP (Acid Phosphatase, Prostate) gene, a known prostate cancer marker gene. Red color denotes higher expression; blue denotes lower expression. The ACPP gene is spatially variable across all three tissue sections. **C.** Mean expression level of all spatially variable glycolysis genes (left) and an example, enolase 1 (ENO1) in section 3.3 **D.** Mean expression level of all spatially variable oxidative phosphorylation genes (left) and an example, NDUFB11 in section 3.3 **E.** Extracellular (SOD3) and mitochondrial (SOD2) superoxide dismutase show spatially distinct expression profiles. SOD2 has the opposite spatial distribution as OXPHOS (S1D).

**Figure S2.**
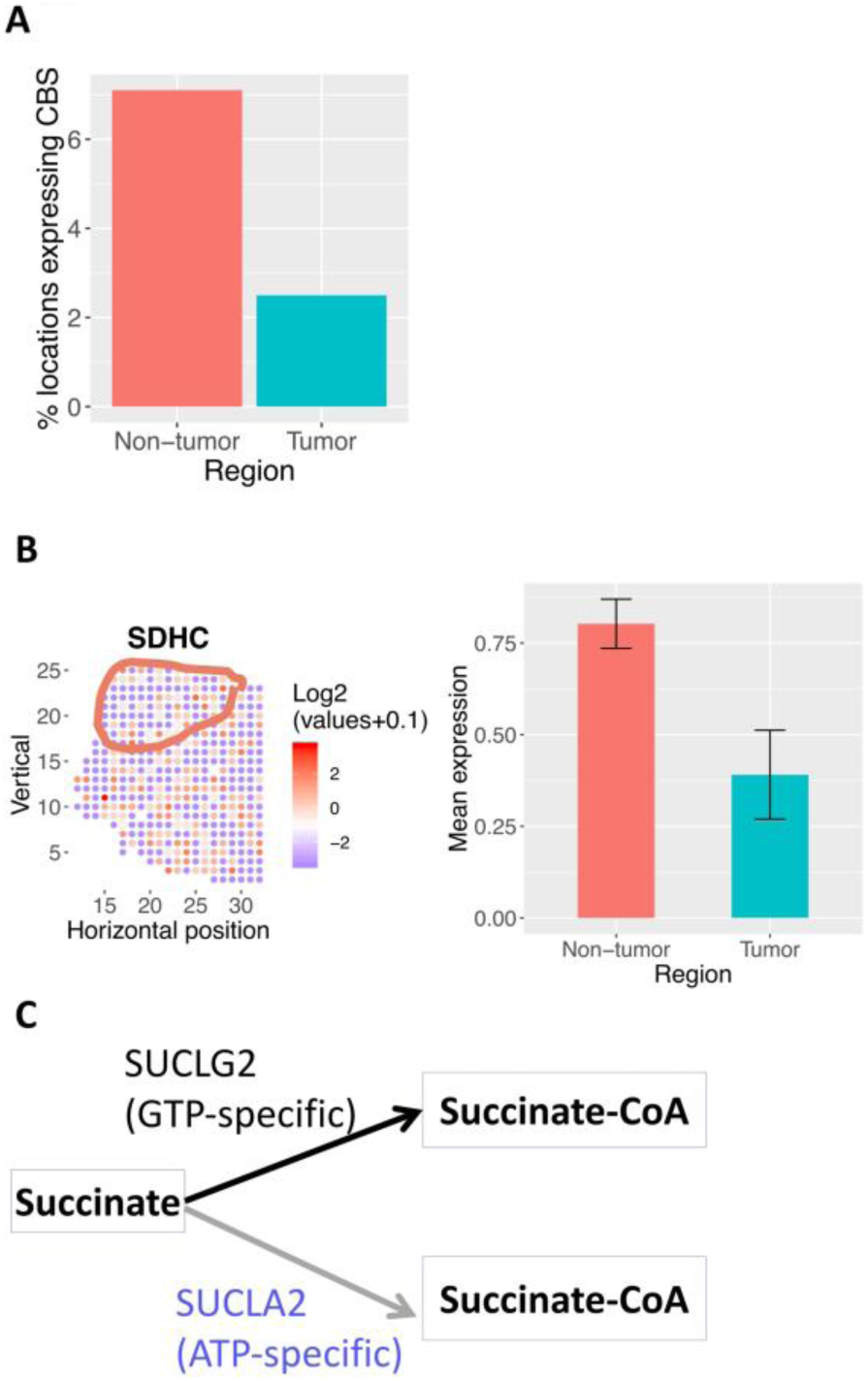
**A.** Bar plot of percentage of locations within tumor and non-tumor regions that express CBS. **B.** Succinate dehydrogenase is depleted in the tumor region of tissue section 1.2. Tumor region is circled. Left: log2 expression of SDHC across the tissue section. Red means higher expression; blue/white means low or no expression. Right: Mean expression of SDHC in non-tumor and tumor region. Error bar represents standard error of the mean. **C.** Model predicted that in tissue section 2.4, SUCLG2 (GTP-specific succinyl-CoA synthetase) is lethal because the alternative route to produce succinate-CoA via SUCLA2 (ATP-specific succinyl-CoA synthetase) is absent in the malignant region. Each rectangle represents a metabolite. Each arrow represents a reaction or transport (black arrow: reaction is present in the tumor; gray arrow: reaction is absent from the tumor). The name of each reaction is labeled above the corresponding arrow.

**Figure S3.**
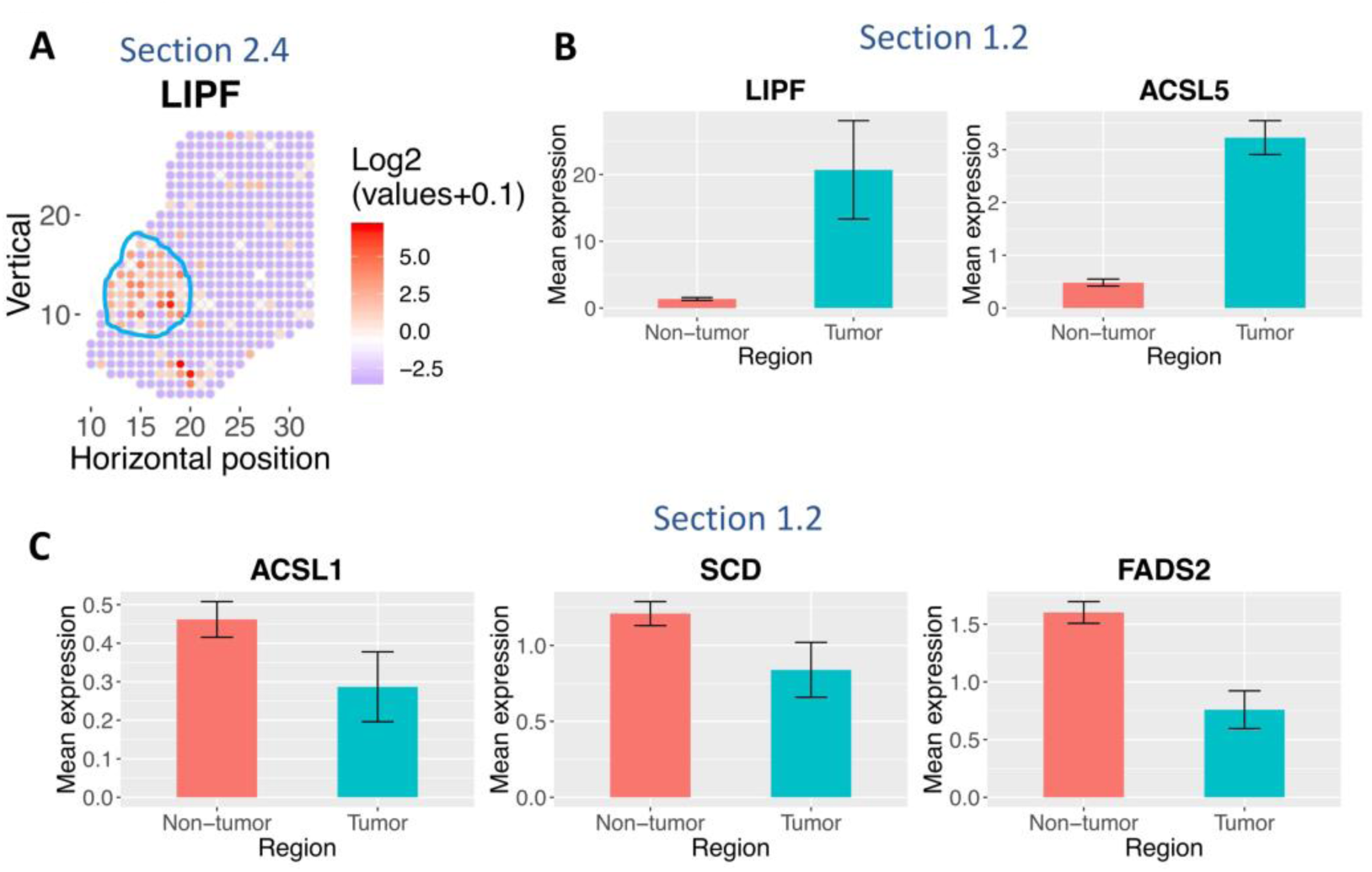
**A.** Lipase F is also enriched in the tumor region in tissue section 2.4. Red color denotes higher expression; blue denotes lower expression. Tumor region is circled by blue lines. **B.** Bar plot of mean expression level of lipolysis gene LIPF and fatty acid synthesis gene ACSL5 in section 1.2.. Error bar represents standard error of the mean. **C.** Bar plot of mean expression level of fatty acid oxidation gene ACSL1, and fatty acid desaturation gene SCD and FADS2 in section 1.2.

**Figure S4.**
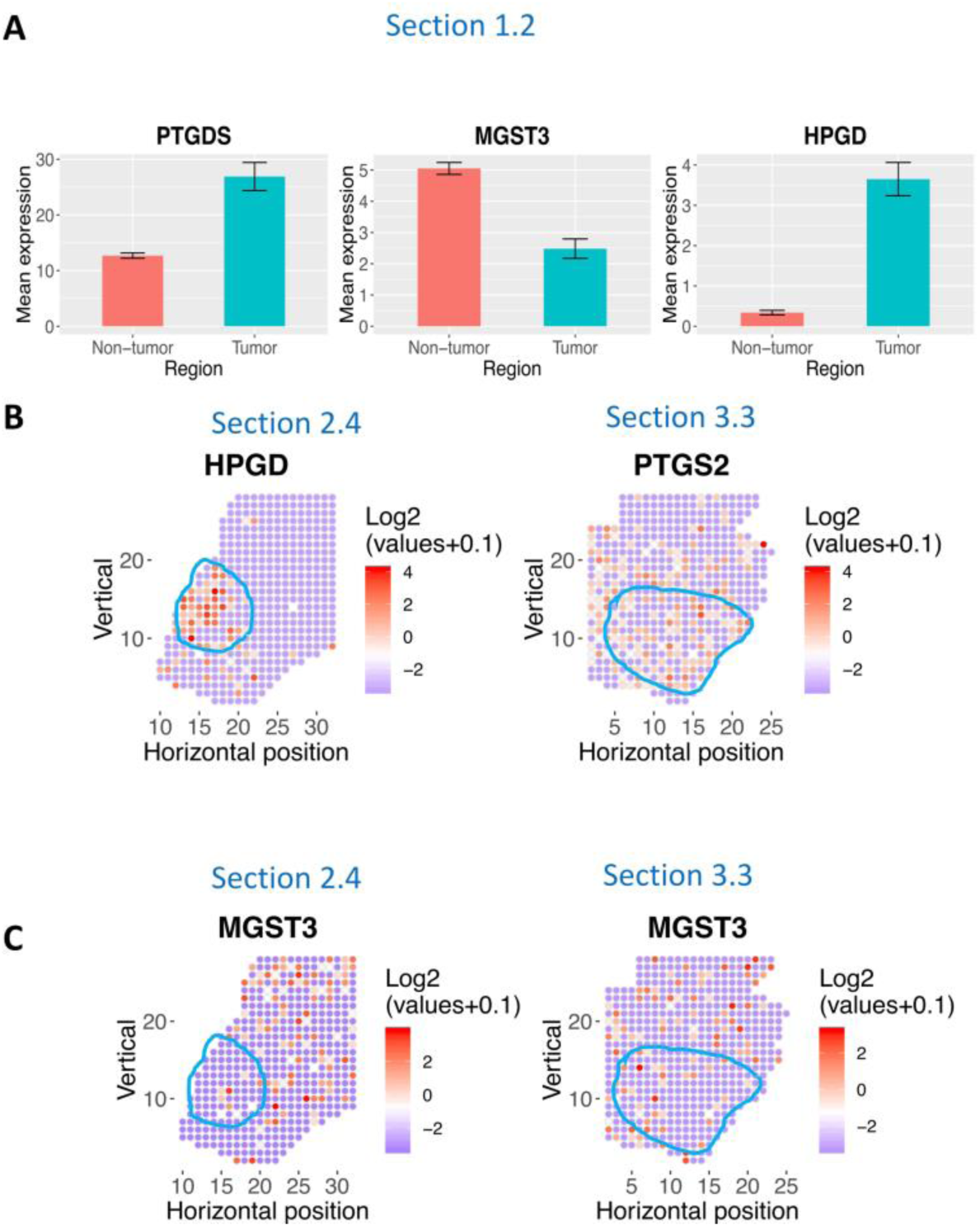
Additional genes in arachidonic acid metabolism are spatially variable in prostate cancer **A.** Bar plot of mean expression level of arachidonic acid metabolism genes in section 1.2. MGST3 is depleted in tumor region while PTGS and HPGD are enriched. Error bar represents standard error of the mean. **B.** HPGD is enriched in the tumor region in tissue section 2.4; PTGS2 (i.e. COX-2), the first step in prostaglandin synthesis, is enriched in tumor region in section 3.3. Red color denotes higher expression; blue denotes lower expression. Tumor regions are highlighted in blue. **C.** MGST3 is spatially variable and depleted in tumor regions in section 2.4 and 3.3 as well. Red color denotes higher expression; blue denotes lower expression. Tumor regions are highlighted in blue.

## Notes

#### Summary of Updates

Updated Abstract, Results & Discussion section

